# Genome-wide analysis of focal DNA hypermethylation in *IDH*-mutant AML samples

**DOI:** 10.1101/2021.03.03.433799

**Authors:** Elisabeth R. Wilson, Nichole M. Helton, Sharon E. Heath, Robert S. Fulton, Jacqueline E. Payton, John S. Welch, Matthew J. Walter, Peter Westervelt, John F. DiPersio, Daniel C. Link, Christopher A. Miller, Timothy J. Ley, David H. Spencer

## Abstract

Recurrent mutations in *IDH1* or *IDH2* in acute myeloid leukemia (AML) are associated with increased DNA methylation, but the genome-wide patterns of this hypermethylation phenotype have not been comprehensively studied in AML samples. We analyzed whole-genome bisulfite sequencing data from 15 primary AML samples with *IDH1* or *IDH2* mutations, which identified ~4,000 focal regions that were uniquely hypermethylated vs. normal CD34+ cells. These regions had modest, but significant, hypermethylation in AMLs with biallelic *TET2* mutations, and 5-hydroxymethylation levels that were dependent on functional TET2, indicating that hypermethylation in these regions is caused by inhibition of TET-mediated demethylation. Focal hypermethylation in *IDH*^mut^ AMLs occurred in regions with low methylation in normal CD34+ cells, implying that DNA methylation and demethylation are active at these loci. AML samples containing *IDH* and *DNMT3A*^R882^ mutations were significantly less hypermethylated, suggesting that methylation in these regions is mediated by DNMT3A. *IDH*^mut^-specific hypermethylation was highly enriched for enhancers that form direct interactions with genes involved in normal hematopoiesis and AML, including *MYC* and *ETV6*. These results suggest that focal hypermethylation in *IDH*-mutant AML occurs by altering the balance between DNA methylation and demethylation, and that disruption of these pathways at enhancers may contribute to AML pathogenesis.

## Introduction

DNA methylation changes in acute myeloid leukemia (AML) occur because of disruptions in the balance between processes that add or remove 5-methyl groups to cytosines^1,2^. In both normal and malignant hematopoietic cells, *de novo* DNA methylation is catalyzed primarily by the DNA methyltransferase DNMT3A^3,4^, which methylates unmethylated DNA substrates. Demethylation occurs both passively after DNA synthesis in the absence of DNMT1-mediated propagation of hemi-methylated DNA, and actively via hydroxylation of 5mC by the TET family of hydroxylases with subsequent removal of modified cytosine residues. Alterations in these opposing forces result in either increased or decreased DNA methylation in AML cells compared to normal hematopoietic cells. These changes include diffuse hypomethylation across large genomic regions, as well as focal hypermethylation in CpG islands (CGIs). We recently showed that CGI hypermethylation in AML is mediated by DNMT3A and is present in nearly all AML subtypes^5^. In addition to these changes, specific DNA methylation patterns correlate with recurrent AML mutations that influence DNA methylation pathways. This includes the *DNMT3A*^R882^ mutation, which impairs DNA methylation activity and results in a focal, canonical hypomethylation phenotype^5^.

Mutations in *IDH1* and *IDH2* are also associated with altered DNA methylation patterns^6,7^ that are thought to occur by disrupting active DNA demethylation. *IDH1* and *IDH2* encode metabolic enzymes not normally involved in DNA methylation, but when mutated produce 2-hydroxyglutarate (2HG)^8^ that inhibits the TET family of enzymes^9^ thereby reducing active demethylation. Analysis of DNA methylation in primary AML samples using array-based technologies and reduced-representation bisulfite sequencing (RBBS) has shown that DNA methylation is increased in specific genomic regions in samples with *IDH* mutations^6,10^. While the direct effects of these changes on gene regulation have been challenging to identify, the contribution of *IDH* mutations to leukemogenesis has been established in mouse models.

Expression of either *IDH1*^R132H^ or *IDH2*^R140Q^ blocks normal hematopoietic differentiation, promotes myeloproliferation^11–13^, and can result in AML transformation in the presence of cooperating mutations^13,14^. These studies demonstrate the contribution of mutations in *IDH1* and *IDH2* to AML development, which may occur by disrupting the balance between DNA methylation and demethylation.

Although previous studies using targeted DNA methylation approaches have established general effects of *IDH1* and *IDH2* mutations on DNA^6,7,10,15^, a genome-wide analysis of methylation patterns in primary AML samples has not yet been described. It is therefore unclear whether *IDH1* vs. *IDH2* mutations manifest unique, mutation-specific methylation phenotypes, and whether these methylation changes are distinct from DNMT3A-mediated CGI hypermethylation. In addition, although *IDH* mutations are thought to cause hypermethylation via inhibition of TET enzymes, the extent of overlap in methylation phenotypes between AML samples with mutations in *IDH1/IDH2* and *TET2* remains unclear. Here, we performed a genome-wide analysis of DNA methylation in primary AML samples with recurrent mutations in *IDH1* or *IDH2* using whole-genome bisulfite sequencing (WGBS). WGBS data from normal hematopoietic cells and AML samples with *TET2* mutations and other mutational profiles were included to define the methylation phenotypes specific to *IDH* mutations, and to determine whether these patterns are present in AML samples with *TET2* mutations. We integrated these data with epigenetic modifications and three-dimensional (3D) genome architecture from primary AML samples to characterize the functional genomic elements that are affected upon disruption of the balance between DNA methylation and demethylation in AML cells.

## Materials and Methods

### Patient samples

Primary AML samples and normal hematopoietic cells for epigenetic studies were obtained from presentation AML and normal bone marrow aspirates, following informed consent using protocol (201011766) approved by the Human Research Protection Office at Washington University as described previously^5^ (Table S1). All experiments with AML samples used total bone marrow cells after estimating the leukemic purity.

### Whole genome bisulfite and oxidative bisulfite sequencing and data analysis

Whole-genome bisulfite sequencing data for 38 samples were described previously^5^. Data for 13 additional samples were generated using 50ng of DNA with the Swift Accel-NGS Methyl-Seq library preparation kit. Oxidative bisulfite sequencing libraries were prepared following treatment of 200ng of DNA with the TrueMethyl oxBS module (Cambridge Epigenetix) prior to bisulfite conversion and Swift library construction and sequencing on NovaSeq 6000 instruments (Table S1). Data were aligned to the GRCh38 human reference and processed into methylated read counts using biscuit^16^ with default parameters. Differentially methylated regions (DMRs) were identified between AML groups using read count data via DSS^17^ and subsequent filtering to retain regions with >10 CpGs and a difference in mean methylation of 0.2. Mutation-specific DMRs were identified using hierarchical clustering where group level mean methylation values at all identified AML DMRs where assessed for euclidean distances via ‘dist’ with default parameters. Clustering topology was analyzed to identify DMRs where a outlier branch represented the specified AML mutational group. Statistical analysis of differential methylation in DMRs was performed on the sum of the methylated/unmethylated counts using a Fisher’s exact test. 5hmC values were obtained by subtracting the methylation ratios from OxBS data from WGBS data at all CpGs with coverage of at least 10x.

### ChIP-seq for histone modifications

ChIP-seq was performed using ChIPmentation^18^ with the following antibodies: H3K27me3 (9733S), and H3K27ac (8173S) from Cell Signaling Technology, and H3K4me1 (ab1012) from Abcam. Sequencing was performed on a NovaSeq 6000 (Illumina, San Diego, CA) to obtain ~50 million 2×150 bp reads. Data were analyzed via adapter trimming with trimgalore and alignment to GRCh38 using bwa mem^19^. Normalized coverage for visualization and analysis used the deeptools “bamCoverage” tool^20^, and peaks were called with MACS2^21^. Statistical comparisons with DESeq2^22^ used raw fragment counts at peak summits, and visualizations were prepared with Gviz^23^. Superenhancer analysis was conducted using ROSE software^24,25^ with default parameters.

### RNA-seq analysis

RNA-seq data from AML samples were obtained from the AML TCGA study^15^. TPM values were obtained using kallisto^26^ and gene counts were generated using the tximport Bioconductor package^27^ in R with the tx2gene option set to accomplish gene-level summarization. Previously published RNA-seq data for normal CD34+ cells generated using the same procedures that were used for the AML samples ^28,29^ were obtained as raw sequencing reads from the short-read archive (GSE48846) and processed as described above.

### Hi-C data analysis

Hi-C data were obtained from previous studies of 3D genome interactions in primary AML samples^30^ and normal hematopoietic stem/progenitors^31^. All libraries were generated using MboI digestion prior to proximity ligation and data were analyzed using the juicer pipeline^32^. Loops were identified with HICCUPs and were analyzed using bedtools^33^ to identify overlap with genes and putative enhancers. Visualizations used the GenomicInteractions and Gviz R packages^23^.

## Results

### Primary AML samples with IDH1 or IDH2 mutations are focally hypermethylated at regions with low methylation in normal hematopoietic cells

We performed WBGS using 15 primary bone marrow aspirate samples from AML patients with canonical *IDH* mutations, including seven with *IDH1*^R132C/G^, seven with *IDH2*^R140Q^, and one with an *IDH2*^R172K^ allele (referred to hereafter as *IDH*^mut^). These data were analyzed with WGBS data from 36 other primary AML samples representing eight mutational categories, including five with biallelic loss-of-function mutations in *TET2,* and primary CD34+ cells from six healthy adult bone marrow donors^5^. All AML samples were previously sequenced using whole genome and/or whole exome sequencing^15,34^, which confirmed that the mutations affecting DNA methylation were present in the dominant leukemic clone at presentation (Figure 1A). Importantly, the *IDH*^mut^ AML samples lacked mutations in *DNMT3A* and *TET2* to minimize effects of other mutations on DNA methylation patterns. WGBS produced a mean of 12x coverage (range: 4-16) for at least 28 million CpGs in the human reference sequence across all samples. *IDH*^mut^ AML samples had genome-wide methylation levels similar to all other samples (Figure 1B), but were more methylated at CGIs compared with normal CD34+ cells and AMLs with *DNMT3A*^R882^ mutations, which have diminished CGI hypermethylation (*IDH*^mut^ vs. CD34+ cells P<0.0001; *IDH*^mut^ vs. *DNMT3A*^R882^ P<0.0001; Figure 1C). CGI methylation levels were similar between *IDH*^mut^ AML samples and AMLs with wild type *IDH1* and *IDH2* (mean CGI methylation of *IDH*^mut^ cases = 0.35, mean of all other groups = 0.33; P= 0.1; Figure 1C), indicating that *IDH* mutations do not result in an exaggerated CGI hypermethylation phenotype.

**Figure 1.**
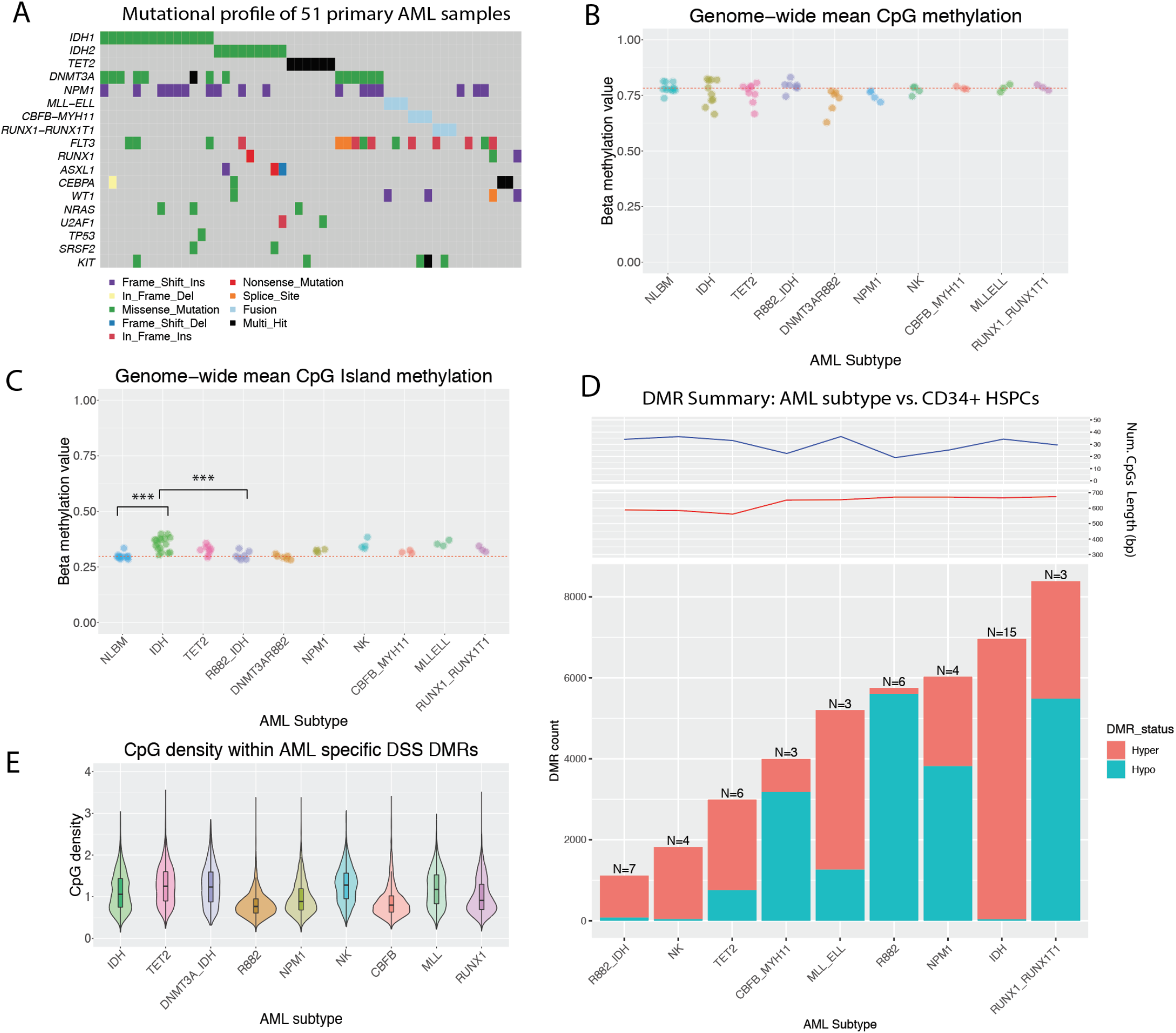
Genome-wide DNA methylation patterns in seven AML mutation groups and normal CD34+ cells. A) Summary of mutation status in 51 primary AML patients with WGBS data. B) Average methylation levels across ~28 million CpGs in CD34+ cells (N=6) and AML subtypes (*IDH1*^mut^ or *IDH2*^mut^, n=15; *TET2*^mut^, n=5; *DNMT3A^R882^,* n=6; *DNMT3A*^R882^/*IDH*^mut^, n=7; normal karyotype with *NPM1c* and wild-type *IDH1, IDH2, TET2,* and *DNMT3A,* n=4; Normal karyotype with wild-type *NPM1, IDH1, IDH2, TET2,* and *DNMT3A,* n=4; *CBFB-MYH11*, n=3; *KMT2A-ELL,* n=3; *RUNX1-RUNX1T1,* n=3). C) Mean CpG island methylation levels in CD34+ cells and AML subtypes. D) Number of differentially methylated regions (DMRs) identified for each AML subtype compared with normal CD34+ cells. Blue and red bars represent hypomethylated and hypermethylated DMRs with respect to normal CD34+ cells, respectively. Mean number of CpGs per DMR (top panel) and DMR length (bottom panel) are shown for each AML subtype. E) Mean CpG density across DMRs identified in each AML subtype.

Differential methylation analysis was performed using established methods^17^ for eight mutation-defined AML groups vs. normal CD34+ cells. This identified a mean of 4,689 (1,114-8,386) Differentially Methylated Regions (DMRs) across the eight categories (minimum methylation difference of 0.2, FDR<0.05); of these, between 6% and 97% were hypermethylated in the AML samples vs. CD34+ cells (see Figure 1D). AMLs with *DNMT3A*^R882^ had the most hypomethylated DMRs (97% of 5,747 DMRs), whereas *IDH*^mut^ AML samples had the most hypermethylated DMRs, with 99% of the 6,960 identified regions having at least 0.2 higher mean methylation than CD34+ cells. *TET2*-mutant samples were also hypermethylated, but at fewer loci, with 74% of the 2,991 identified DMRs having higher methylation. The fewest DMRs were identified in samples with *IDH1* or *IDH2* mutations and *DNMT3A*^R882^, which is consistent with previous studies suggesting that AML samples with both mutations have abrogated methylation phenotypes^10^. Analysis of DMRs identified in *IDH*^mut^ samples demonstrated that 29% were associated with promoters and 43% occurred in CGIs, which was similar to the frequencies observed for commonly hypermethylated DMRs in other AML subtypes (Figure S1A). Interestingly, although *IDH* mutations are thought to lead to hypermethylation by inhibiting active demethylation, most *IDH*^mut^-specific DMRs had low methylation in normal hematopoietic cells. For example, 72% of the *IDH*^mut^ DMRs had a mean methylation <0.3 in both CD34+ cells (Figure S1B) and more mature myeloid cell populations (Figure S1C), suggesting that DNA methylation pathways must be active in these regions to achieve the level of hypermethylated observed in *IDH*^mut^ AML samples.

### IDH^mut^-specific methylation changes are distinct from AML-associated CGI hypermethylation and are influenced by IDH mutation type

Because AML patients with *IDH1* vs. *IDH2* mutations have distinct clinical and molecular phenotypes^35–39^ we next sought to identify methylation changes uniquely associated with *IDH1* and/or *IDH2* mutations and to determine whether these patterns are distinct from AML-associated CGI hypermethylation (Figure 2A). Thus, we performed hierarchical clustering of mean methylation values for either *IDH1*^mut^ or *IDH2*^mut^ and *IDH*^wt^ AML samples at DMRs identified above, and used the clustering topology to identify regions where the single outlier branch represented the *IDH* mutant AML samples (Figure 2B, see Methods). Samples with *TET2* and/or *DNMT3A*^R882^ mutations were excluded, since they may share methylation changes with *IDH*^mut^ samples, or lack CGI hypermethylation^5,6^, respectively. This approach identified 3,928 *IDH1*^mut^-specific and 1,821 *IDH2*^mut^-specific DMRs, of which 90% and 79% were hypermethylated with respect to normal CD34+ cells, respectively (see Figure 2C-E). Consistent with the analysis above, most DMRs displayed low methylation in normal cells, with 55% of *IDH1*^mut^-specific and 71% of *IDH2*^mut^-specific loci having a methylation level <0.3 in CD34+ cells (Figure 2C-E) and mature myeloid cells (Figure S1D). There was extensive overlap between the *IDH* mutation-specific DMRs (94%, 5,403 of 5,749 total DMRs), and AML samples with either mutation were hypermethylated at both DMR sets (Figure 2E). However, hierarchical clustering demonstrated considerable variability in methylation levels between the *IDH1*^mut^ and *IDH2*^mut^ samples (Figure 2F). This was most striking for the *IDH2*^mut^ samples, which included three AMLs with lower methylation at the *IDH2*^mut^-specific DMRs (Figure 2F-G). *IDH2*^mut^ AML samples were less methylated at the combined set of *IDH1*^mut^-specific and *IDH2*^mut^-specific DMR loci (*IDH2*^mut^= 0.54 vs. *IDH1*^mut^=0.70; p-value= 0.04), although they remained hypermethylated relative to CD34+ cells (Figure 2G). This difference was not related to mutant *IDH* allele abundance (all samples had VAFs >30%, Table S1), and did not correlate with other recurrent mutations, including *NPM1c* (4 in *IDH1*^mut^ and 3 in *IDH2*^mut^ samples, Figure 2F).

**Figure 2.**
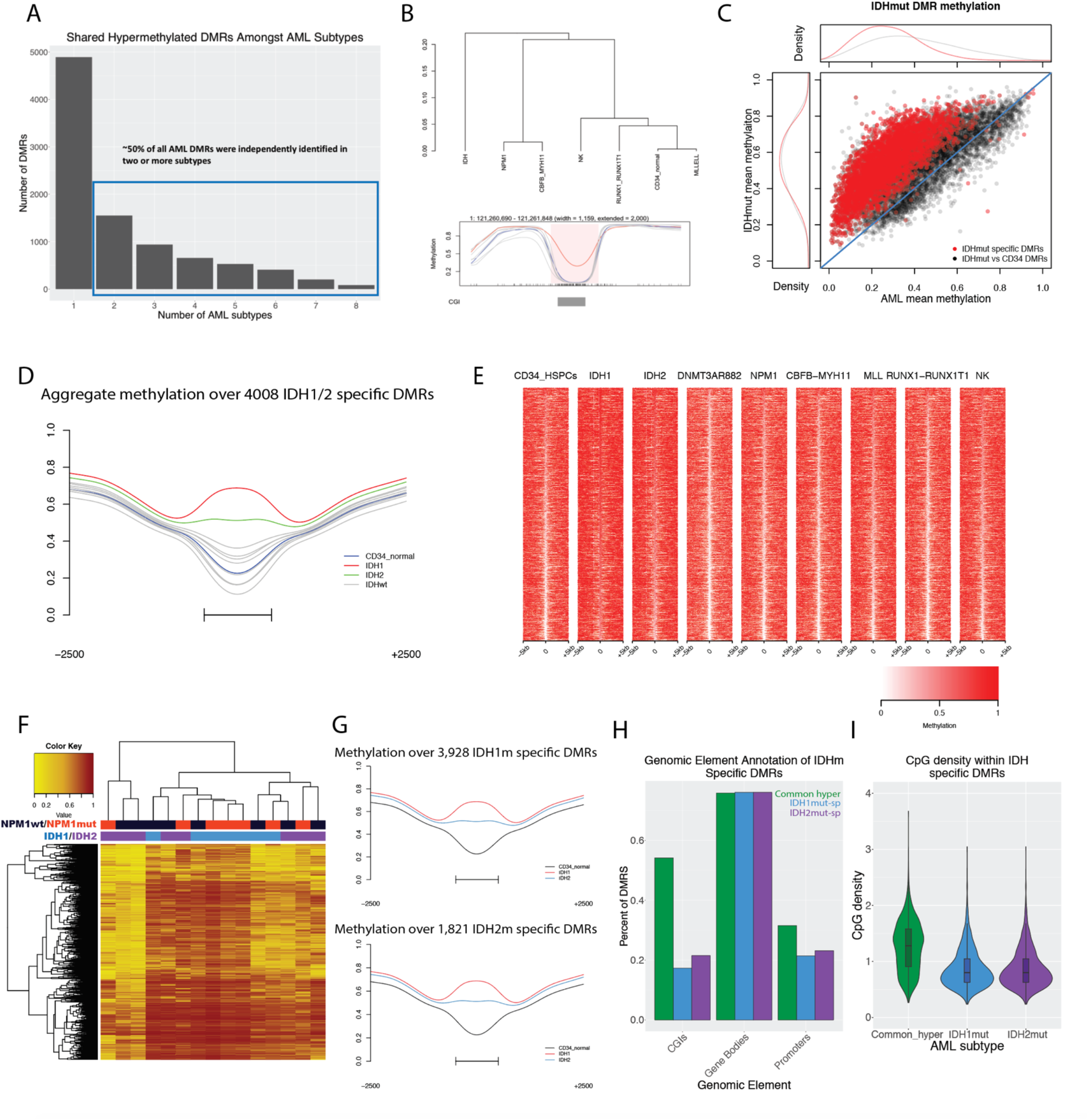
Characterization of *IDH*^mut^-specific DMRs. A) Distribution of shared hypermethylated DMRs across AML subtypes. B) Example of the hierarchical clustering approach used to identify *IDH*^mut^-specific DMRs. Aggregate DMR methylation for individual AML subtypes are shown in bottom panel with *IDH* samples indicated in red and CD34+ cells indicated in blue. C) Mean DMR methylation across *IDH*^mut^-specific DMRs in *IDH*^mut^ samples versus mean methylation of all other AMLs. D) Aggregate methylation levels across all *IDH1/2*^mut^-specific DMRs in *IDH*^mut^ and *IDH*^wt^ AML subtypes. E) Locus heatmap showing mean methylation values by group for all *IDH*^mut^-specific DMRs, where each column is centered over the DMR with the window extending 5kb up- or down-stream the DMR center. F) Mean DMR methylation across all *IDH1/2*^mut^-specific DMRs (rows) in 15 individual *IDH*^mut^ cases (columns). G) Aggregate DMR methylation across 3,928 *IDH1*^mut^ specific DMRs and 1,821 *IDH2*^mut^ specific DMRs respectively. H) Fraction of functional genomic elements overlapping generically hypermethylated DMRs, *IDH1*^mut^ specific DMRs, and *IDH2*^mut^ specific DMRs. I) Distribution of CpG densities across generically hypermethylated regions in primary AML and *IDH1*^mut^- and *IDH2*^mut^-specific DMRs.

To determine whether *IDH*^mut^-specific hypermethylation is similar to the canonical CGI hypermethylation seen in other AML samples, we analyzed the CpG content and genomic features of the *IDH*^mut^-specific DMRs. Interestingly, the CpG density and overlap with genomic annotations was markedly different for both *IDH1*^mut^-specific and *IDH2*^mut^-specific DMRs compared to regions with AML-associated CGI hypermethylation. For example, the combined set of *IDH*^mut^-specific DMRs displayed significantly lower CpG density (mean CpG density of 0.89 vs. 1.26; P<0.0001) and less overlap with annotated CGIs (23% vs. 54%) compared to 4,573 commonly hypermethylated regions identified in at least 2 other AML mutation categories (Figure 2H-I). Promoter regions were also underrepresented among the *IDH*^mut^-specific DMRs (21% vs 31%; see Figure 2H), consistent with depletion of promoter-associated CGIs among *IDH*^mut^-specific DMRs, further suggesting that *IDH*-associated hypermethylation is distinct from AML-associated CGI hypermethylation.

### Hypermethylation in TET2^mut^ AMLs overlaps with IDH^mut^-specific hypermethylation, but does not phenocopy the extent of methylation changes

We next determined whether AML samples with biallelic loss-of-function mutations in *TET2* shared similar genome-wide patterns of hypermethylation with *IDH*^mut^ AMLs. Initial comparison of the *TET2*^mut^ AMLs vs. normal CD34+ cells yielded fewer DMRs and a lower proportion of hypermethylated regions compared to *IDH*^mut^ samples (2,512 vs. 6,523 DMRs, and 78% vs 99% hypermethylated regions, respectively), consistent with previous reports that inactivation of *TET2* has a less profound effect on DNA methylation^6,10^. Identification of *TET2*^mut^-specific DMRs using the clustering approach described above (with *IDH*^mut^ and *DNMT3A*^R882^ AMLs excluded from the analysis) produced only 51 *TET2*^mut^-specific DMRs, confirming that these AMLs lack a strong hypermethylation phenotype (Figure 3A). Although many DMRs were hypermethylated relative to CD34+ cells (31 of 51), the fraction was significantly less than in either *IDH1*^mut^ or *IDH2*^mut^ AMLs (60% vs 91% and 73%, respectively). *TET2*^mut^-specific DMRs were also not enriched for either CGIs or promoters, compared to the set of commonly hypermethylated DMRs (17% of *TET2* DMRs vs. 54% generic DMRs; 11% of *TET2* DMRs vs. 31% generic DMRs; see Figure S2A-B), suggesting these regions are unlikely to reflect CGI hypermethylation.

**Figure 3.**
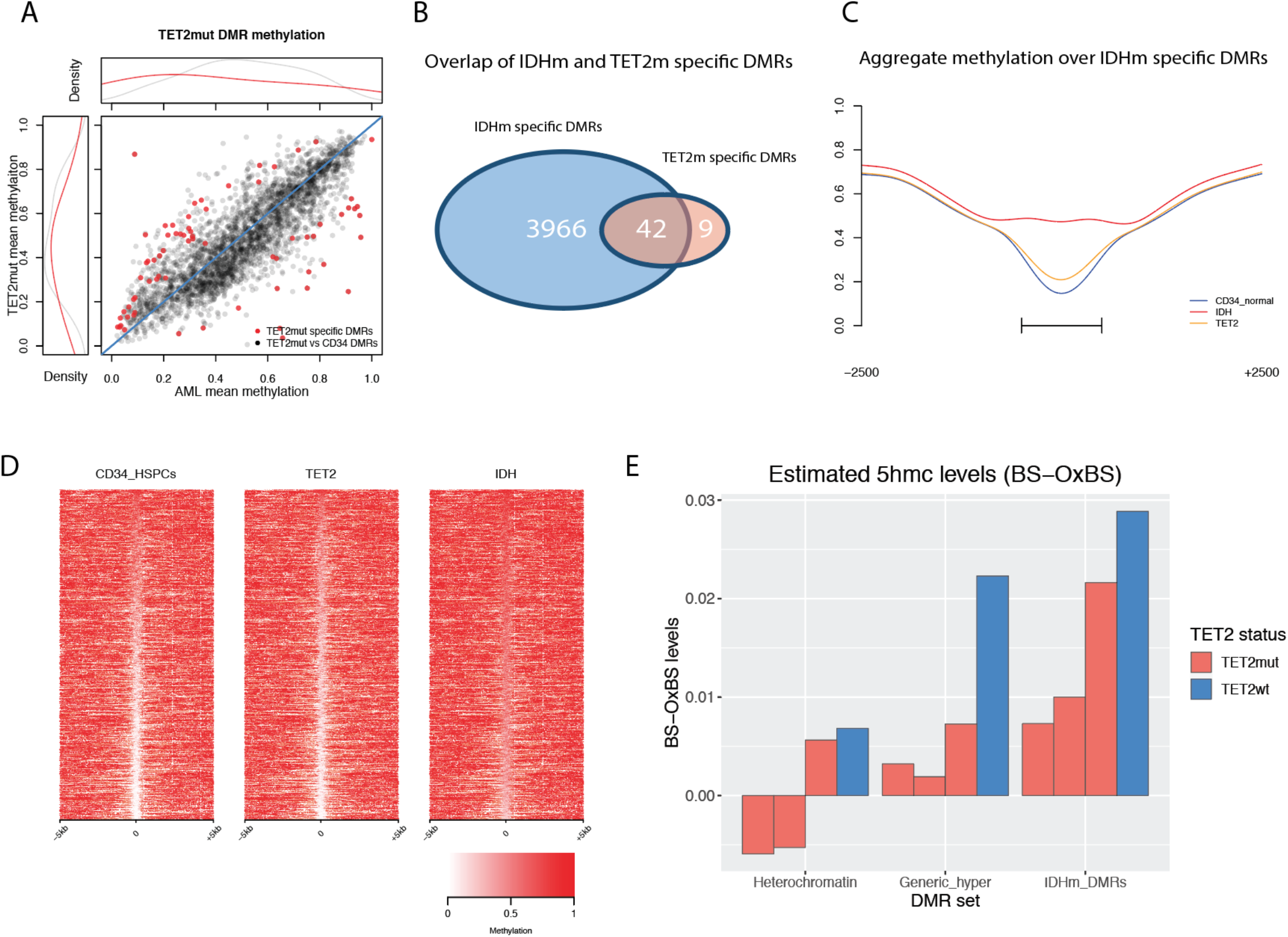
*TET2*^mut^ AMLs have modest hypermethylation that overlaps *IDH*^mut^-specific DMRs. A) Mean DMR methylation across 2512 in *TET2*^mut^ DMRs called vs. CD34+ cells (black points) and 52 in *TET2*^mut^-specific DMRs (red points) in *TET2* mutant samples versus mean methylation in all other AMLs. B) Intersection of in *TET2*^mut^-specific and *IDH*^mut^-specific DMRs. C) Aggregate methylation over *IDH*^mut^-specific DMRs in *IDH*^mut^ and in *TET2*^mut^ AML and CD34+ cells. D) Locus heatmap of mean methylation values for all *IDH*^mut^-specific DMRs (rows), where each column is centered over the DMR with the window extending 5kb up- and down-stream the DMR center point. E) Mean 5hmC (WGBS minus oxWGBS) levels in in *TET2*^mut^ and in *TET2*^wt^ AML samples at 4650 ChromHMM heterochromatic regions, 4586 generically hypermethylated regions, and 4008 *IDH*^mut^-specific hypermethylated DMRs.

To investigate the interaction between *IDH* mutations and *TET2*-mediated demethylation, we compared *TET2*^mut^-specific and *IDH*^mut^-specific DMRs, and performed oxidative bisulfite sequencing^40^ to measure 5-hydroxymethylation (5hmC) in samples with and without biallelic inactivating *TET2* mutations. This analysis showed that 82% (42 of 51) of the *TET2*^mut^-specific DMRs overlapped an *IDH*^mut^-specific hypermethylated region (Figure 3B). Methylation in *TET2*^mut^ AMLs at the 4008 *IDH*^mut^-specific DMRs was also significantly increased compared to CD34+ cells (mean methylation of 0.38 vs. 0.31; 61% of DMRs with increased methylation via Fisher’s exact test with q<0.05; Figure 3C-D). We next analyzed levels of 5hmC at *IDH*^mut^ DMRs using paired oxidative and standard whole-genome bisulfite sequencing (oxWGBS and WGBS) of primary AML samples with (N=3) and without (N=1) *TET2* mutations (Figure S2C). Subtraction of oxWGBS from WGBS data for CpGs with >10x coverage demonstrated low levels of 5hmC across the genomes of these samples (mean: 0.51-0.71% in *TET2*^mut^, 1.35% in *TET2*^wt^; Figure S2D), and identifiable peaks at selected loci (Figure S2E). *IDH*^mut^ DMRs had higher levels of 5hmC than in either constitutively methylated heterochromatin or regions with CGI hypermethylation, and 5hmC levels were lower in all three of these regions in *TET2*^mut^ AML samples (Figure 3E, Figure S2F), indicating that 5hmC in these regions is dependent on TET2 activity.

### DNA hypermethylation in IDH^mut^ AML cells requires DNMT3A

To assess whether *de novo* DNA methylation by DNMT3A contributes to *IDH*^mut^-associated hypermethylation, we analyzed methylation levels at *IDH*^mut^-specific DMRs in seven AML samples with co-occurring *IDH1* (N=5) or *IDH2* (N=2) and *DNMT3A*^R882^ mutations, which have a dominant negative phenotype and a more severe hypomethylation than other *DNMT3A* mutations^4^. Interestingly, although *DNMT3A*^R882^/*IDH*^mut^ AMLs were still hypermethylated at *IDH*^mut^-specific DMRs, the degree of hypermethylation was diminished, with 51% of these regions having significantly lower DNA methylation levels than samples with *IDH* mutations alone (Figures 4A-C, S3A). Similar findings were observed in 7 additional *DNMT3A*^R882^/*IDH*^mut^ AML samples using methylation array data from the TCGA AML study^15^ (Figure S3B). To further characterize the extent of this interaction at regions known to be methylated by DNMT3A, we analyzed DNA methylation levels in *DNMT3A*^R882^/*IDH*^mut^ AML samples at hypomethylated DMRs in AMLs with the *DNMT3A*^R882^ allele^5^. Surprisingly, these regions remained nearly fully methylated in the *DNMT3A*^R882^/*IDH*^mut^ double mutant samples, with 93% of the regions having significantly higher methylation than AMLs with *DNMT3A*^R882^ alone (Figures 4D-F, Figure S3C). Similar findings were observed in the array-based methylation data from the TCGA AML study^15^ (Figure S3D), further implying that DNMT3A-mediated methylation and TET-mediated demethylation occur at the same places in the genome.

**Figure 4.**
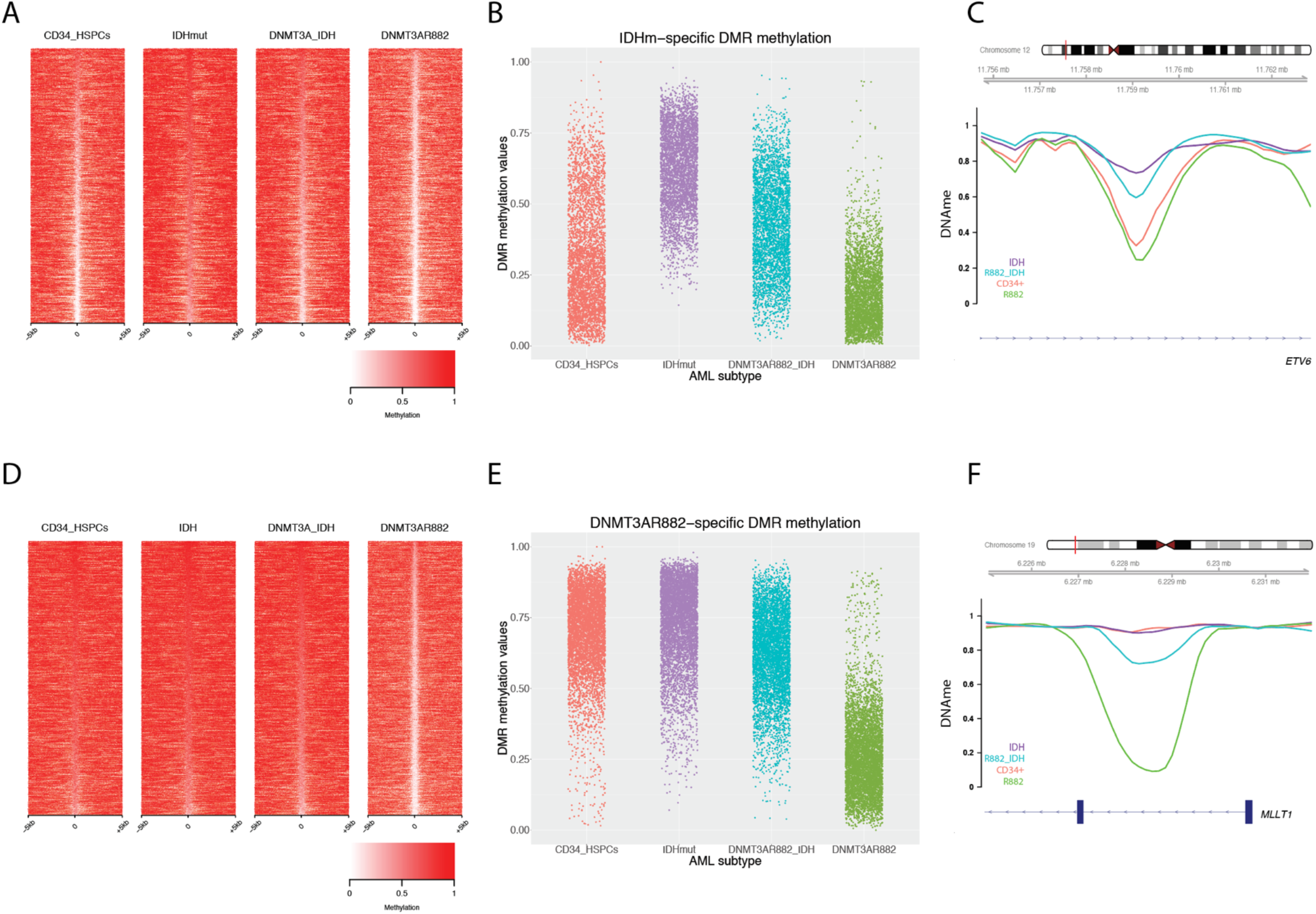
*DNMT3A*^R882^/*IDH*^mut^ double mutant AMLs display an attenuated focal hypermethylation phenotype. A) Locus heatmap of mean methylation at *IDH*^mut^ DMRs (rows) in *IDH1* or *IDH2* mutant, *DNMT3A*^R882^/*IDH*^mut^ double mutant, and *DNMT3A*^R882^ AMLs, and CD34+ cells. B) Distribution of *IDH*^mut^-specific DMR methylation levels by AML subtype. C) Example *IDH*^mut^-specific DMR locus within the *ETV6* gene demonstrating an intermediate methylation phenotype of double mutant samples with respect to *IDH*^mut^ and *DNMT3A*^R882^ mutant AMLs. D) Methylation locus heatmap of average subtype methylation across *DNMT3A*^R882^ DMRs called vs. CD34+ cells in *IDH*^mut^, *DNMT3A*^R882^/*IDH*^mut^ double mutant, and *DNMT3A*^R882^ AMLs, and CD34+ cells. E) Distribution of *DNMT3A*^R882^ DMR methylation levels by AML subtype. F) Example *DNMT3A*^R882^ DMR locus within the *MLLT1* gene, demonstrating the hypomethylation phenotype of *DNMT3A*^R882^ mutant samples with respect to *IDH*^mut^ and *DNMT3A*^R882^/*IDH*^mut^ double mutant AML samples.

### IDH^mut^-specific hypermethylated DMRs are enriched for enhancers

We next asked whether these *IDH*^mut^-specific DMRs were associated with certain chromatin states. Annotation of these DMRs with chromatin states in CD34+ cells^41^ demonstrated that 44% occurred the enhancer chromatin state, which was a 2-fold enrichment over regions commonly hypermethylated (Figure 5A). Similar analysis performed on DMRs identified in other AML subtypes showed a different distribution of chromatin states (Figure S4A-C). We further defined this association using ChIP-seq peaks for enhancer-associated H3K27ac and H3K4me1 modifications from 16 primary AML samples, including 14 *IDH*^wt^ and 2 *IDH*^mut^. Analysis defined 44,762 and 6,917 consensus peaks for H3K27ac and H3K4me1, respectively, which along with H3K27me3 ChIP-seq data from 9 AML patients were used to identify active, poised, and weak enhancer loci. Analysis of the *IDH*^mut^ DMRs showed that 47% overlapped an active enhancer, compared to 3% and 1% that overlapped poised and weak regions, respectively (Figures 5B-D). In comparison, commonly hypermethylated regions showed less overlap with active enhancers (13% of DMRs) and greater intersection with repressive H3K27me3 marks (Figure 5D). We analyzed *IDH*^mut^-specific DMRs for transcription factor (TF) binding motifs, which identified binding sites for hematopoietic-associated TFs, including *SPI1, RUNX1,* and *MYC* (Figure 5E), further supporting the occurrence of *IDH*^mut^-specific hypermethylation at regions with potential regulatory activity. However, quantitative analysis of H3K27ac signal over these regions in samples with and without *IDH* mutations did not identify appreciable differences in wild type vs. mutant samples (p-value=0.24, Figure 5F), suggesting that hypermethylation does not modify H3K27ac levels within these regions.

**Figure 5.**
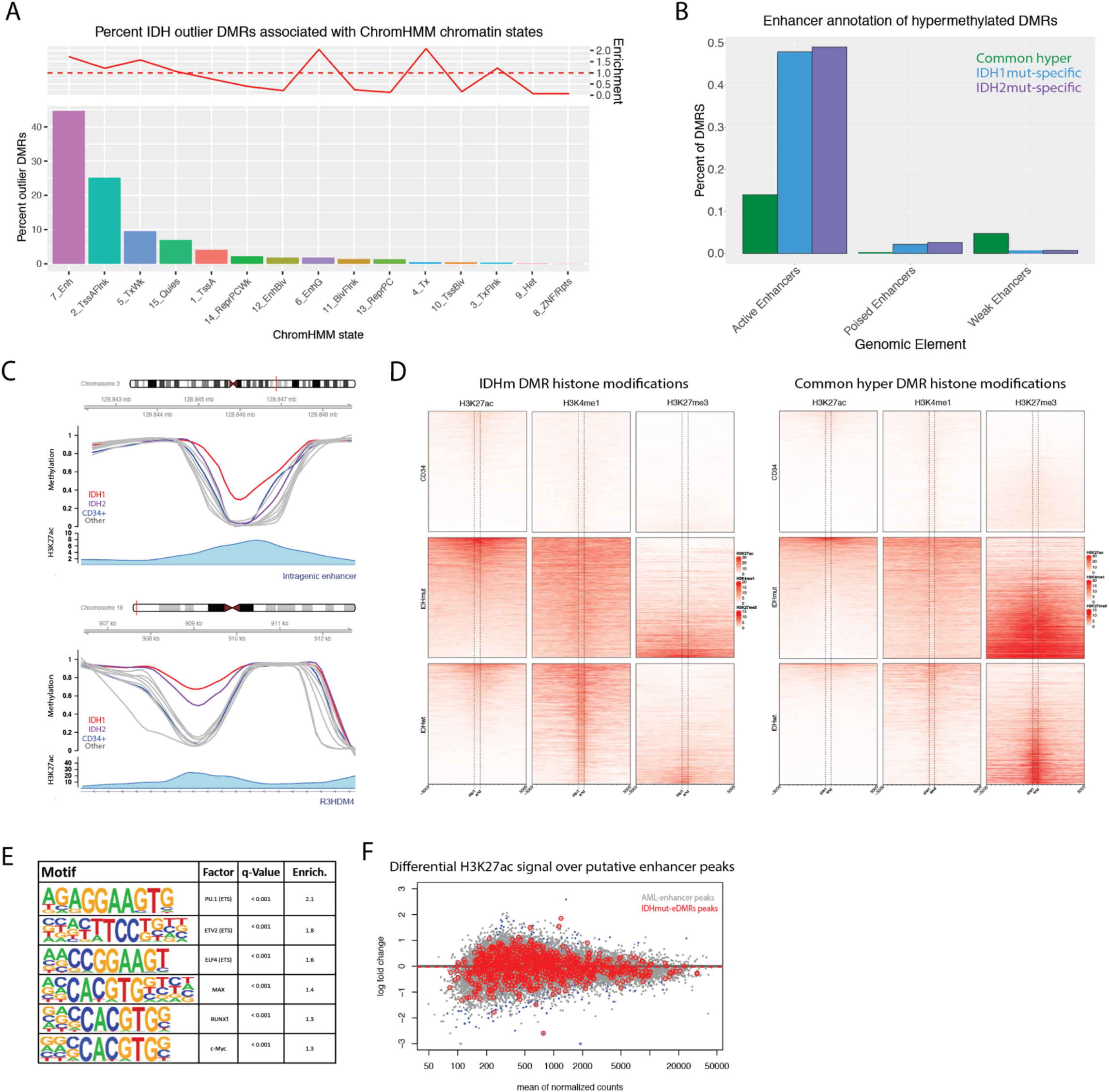
*IDH*^mut^-specific DMRs are enriched for putative enhancers. A) Distribution of ChromHMM chromatin states from CD34+ cells represented in *IDH*^mut^-specific DMRs. Enrichment of chromatin states within *IDH*^mut^-specific DMRs is shown with respect to the frequency of states overlapping regions of common CpG island hypermethylation. B) Enhancer-based annotation of common hypermethylated regions, *IDH1*^mut^, and *IDH2*^mut^ DMRs, where DMRs intersecting an H3K27ac peak alone or in combination with H3K4me1 constitute active enhancers, an H3K27ac peak in combination with H3K27me3 constitutes poised enhancers, and H3K4me1 alone constitutes a weak enhancer profile. C) Examples of intragenic and genic enhancer regions exhibiting *IDH1*^mut^, *IDH2*^mut^, or *IDH1/2*^mut^ hypermethylation compared with CD34+ cells and other AML subtypes. D) Heatmap of enhancer histone modifications and heterochromatin modifications over *IDH*^mut^-specific DMRs (left) and generic hypermethylation (right) in CD34+ cells (N=4 H3K27ac, N=7 H3K3me1, and N=7 H3K27me3), *IDH*^mut^ AML (n=3), and *IDH*^wt^ AML samples (N=9 H3K27ac, N=10 H3K3me1, and N=24 H3K27me3). E) Differential active enhancer signal (H3K27ac) for all AML-associated putative enhancers (black points) compared to putative enhancers intersecting an *IDH1/2*^mut^-specific DMR (red points). F) HOMER motif enrichment analysis of *IDH1/2*^mut^-specific DMRs with respect to a background set of generically hypermethylated regions. G) Enrichment analysis of TF binding events for 445 TFs within *IDH1/2*^mut^-specific DMRs.

### IDH^mut^-specific DMRs occur in enhancers that form direct interactions with highly expressed genes in AML cells

We next asked whether enhancers with *IDH*^mut^ DMRs could be involved in controlling the expression of genes relevant for AML pathogenesis. To directly link these enhancers to their target genes, we analyzed three-dimensional (3D) genome interactions generated using *in situ* HiC from both normal CD34+ cells^31^ and 3 primary AML samples^30^ (all wild-type for *DNMT3A, IDH1, IDH2,* and *TET2)*. This analysis demonstrated that 25% (1047/4008) of all *IDH*^mut^-specific DMRs and 32% (322/1021) of the DMRs in putative enhancers overlapped the ‘loop anchor’ of a genome interaction (Figure 6A, Figure S5A). *IDH*^mut^ DMRs in these loop anchors were highly enriched in ‘superenhancers’, with between 37 and 39% of superenhancers defined in 3 primary *IDH*^mut^ AML samples containing at least one *IDH*^mut^-specific DMR (Figure 6B-C, Figure S5B-C).

**Figure 6.**
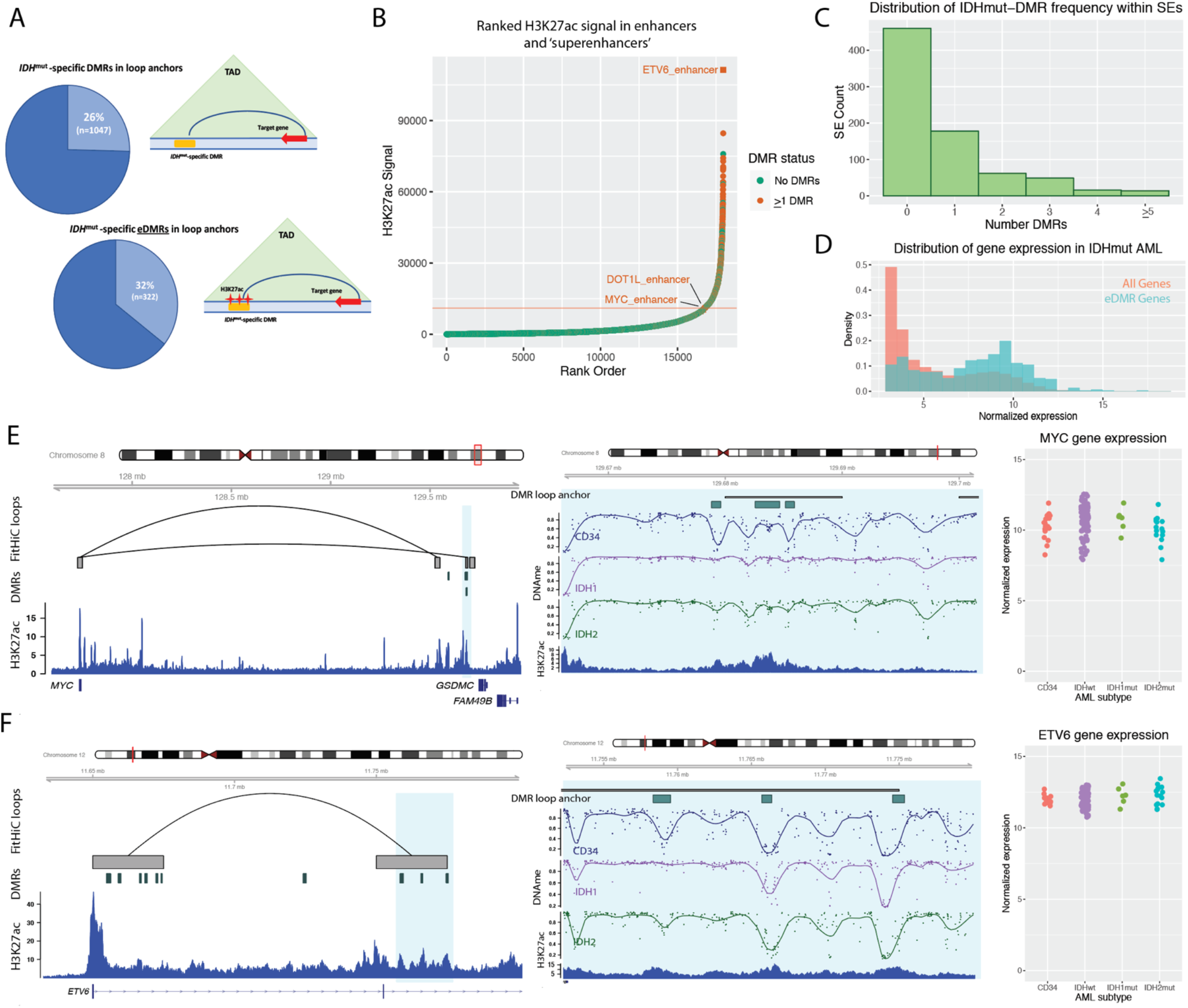
*IDH*^mut^-specific DMRs are enriched in superenhancers and interact with highly expressed genes in AML. A) Schematic of DMR and enhancer-associated DMRs (eDMR) and their interaction with target genes based on intersection HiC-defined genome loops. B) Rank ordered enhancer regions based on H3K27ac signal in a representative *IDH*^mut^ AML sample, annotated by presence of overlapping *IDH*^mut^-specific DMRs (absence of DMRs indicated by green points, greater than one DMR indicated by orange points) and computationally-defined ‘superenhancer’ (above red line). Enhancers of specific hematopoietic genes are designated. C) Distribution of number of *IDH*^mut^-specific DMRs overlapping a set of AML consensus superenhancers (SEs) from H3K27ac data from 4 primary samples (N=779). D) Distribution of normalized gene expression values for all expressed genes (orange histogram) and a set of 750 eDMR target genes (blue histogram) in *IDH*^mut^ AML samples. E) Example *IDH*^mut^ -eDMR locus displaying interactions with the *MYC* promoter. A zoomed in view of the locus demonstrates focal enhancer hypermethylation in *IDH1*^mut^ (purple) and *IDH2*^mut^ (green) samples compared with CD34+ cells (blue). Normalized *MYC* expression is shown for 17 CD34+ cord blood cell samples, 6 and 14 *IDH1*^mut^ and *IDH2*^mut^ samples, and 91 *IDH*^wt^ samples. F) Example *IDH*^mut^-DMR locus in a candidate enhancer that displays robust interactions with the *ETV6* promoter. A zoomed in locus view demonstrates focal enhancer hypermethylation in *IDH1*^mut^ (purple) and *IDH2*^mut^ (green) samples compared with CD34+ cells (blue). Normalized *ETV6* expression is shown for CD34+ cells, *IDH1*^mut^ and *IDH2*^mut^ samples, and *IDH*^wt^ samples (see E for sample sizes).

We next analyzed gene expression in 750 genes with promoters that formed 3D interactions with *IDH*^mut^-specific DMRs using RNA-seq data from 179 AML samples from the TCGA AML study. This showed that the genes linked to *IDH*^mut^-specific DMRs were highly expressed, with 68% of these genes ranked in the top 25^th^ percentile of gene expression (Figure 6D, Figure S5D). Further analysis of 3D genome interactions containing *IDH*^mut^-specific DMRs identified known and novel enhancers of genes important in hematopoiesis and AML, including an enhancer of *MYC*^42–44^ (Figure 6E), and previously unreported putative enhancers that form interactions with *ETV6* (Figure 6F), *DOT1L,* and *SRSF3* (Figure S5E-F). Although we did not observe significant changes in expression of these genes between *IDH*^mut^ and *IDH*^wt^ AMLs, their high expression in AML samples and CD34+ cells was consistent with the enrichment of *IDH*^mut^-specific DMRs in enhancers of active genes (Figure 6D-F).

## Discussion

Recurrent gain-of-function *IDH1* and *IDH2* mutations are known to increase DNA methylation, though the extent and functional consequences of these changes have not been clearly defined. We used whole-genome bisulfite sequencing of primary AML samples to demonstrate that the effects of *IDH* mutations on methylation do not manifest as diffuse changes across the genome, but instead occur in thousands of focal regions that are uniquely hypermethylated compared to both normal CD34+ cells and other AML samples. These regions had lower CpG density and fewer CGIs than loci that are commonly hypermethylated in AML, suggesting that mechanisms of *IDH*^mut^-associated methylation changes are distinct from ‘traditional’ CGI hypermethylation associated with AML. The *IDH2*-mutant AML samples in our dataset had less pronounced hypermethylation than those with *IDH1* mutations, but both were hypermethylated at a highly overlapping set of loci. AMLs with biallelic inactivating mutations in *TET2* had a far less dramatic methylation phenotype, though many *IDH*^mut^-specific DMRs were significantly hypermethylated in *TET2*^mut^ AML samples. Further, oxidative bisulfite sequencing demonstrated increased levels of 5hmC in these regions, which was absent in the *TET2*^mut^ AML samples, providing additional evidence in primary AML samples that both *IDH* and *TET2* mutations cause increased DNA methylation by impairing the TET-mediated DNA demethylation pathway. Regions with *IDH*^mut^-specific hypermethylation were enriched for active enhancers, many of which formed direct interactions with genes that are highly expressed in AML cells, including regulatory sequences that interact with the promoters of *MYC* and *ETV6*. Although the increased methylation at these loci was not associated with repressed chromatin or lower gene expression in *IDH*^mut^ AML samples, this finding demonstrates that *IDH*^mut^-associated hypermethylation affects the regulatory sequences of genes that may contribute to AML pathogenesis.

This study adds new context to the dynamics of *de novo* DNA methylation and active demethylation pathways in normal hematopoietic cells and in AML. The fact that *IDH*^mut^-associated hypermethylation occurs at regions with low levels of DNA methylation in normal CD34+ cells means that *de novo* DNA methylation and TET-mediated demethylation can both be active in these regions, despite their low steady-state methylation levels. This is supported by the observation that AML samples with co-occurring *IDH* and *DNMT3A*^R882^ mutations show significantly attenuated hypermethylation, and that there are appreciable levels of 5hmC in *IDH*^mut^-specific DMRs, which is produced from 5mC as a substrate. Remodeling of DNA methylation by these processes in specific regions has been reported previously in studies of embryonic stem cells, which have shown that methylation and active demethylation are in equilibrium at many loci^1,2^, and may be maintained by the occupancy of methylation and demethylation complexes^45^. Our analysis suggests this equilibrium exists in normal CD34+ cells and is disrupted in the presence of mutant *IDH* alleles, leaving *de novo* DNA methylation unopposed. The focal nature of *IDH*^mut^-associated hypermethylation implies that activity (or occupancy) of DNMT3A and TET enzymes may be ‘concentrated’ in specific genomic regions. The genomic or epigenetic features directing this activity remain unclear^46^, but the enrichment of *IDH*^mut^ DMRs in active enhancers suggests that components of active chromatin may recruit methylation and demethylation machinery. The convergence of these processes at enhancers could provide clues as to why mutations with opposite effects on DNA methylation both contribute to AML development via dysregulation of common target genes.

Our analysis of 3D genome interactions involving *IDH*^mut^-specific DMRs found that these sequences directly interact with genes that are highly expressed in hematopoiesis and AML (e.g., *MYC* and *ETV6*). Contrary to the canonical relationship between DNA methylation and activity, hypermethylation in the *IDH*^mut^ AML samples does not appear to repress either the enhancer elements or expression of their target genes. Other regulatory factors may therefore be dominant to DNA methylation at these loci, and result in persistently high gene expression. It could also be that regions of active chromatin, such as enhancers (and superenhancers), have high rates of methylation turnover and are therefore susceptible to perturbations in methylation and demethylation^1,2^. Focal hypermethylation may occur in DNA elements bound by factors that contribute to ‘fine-tuning’ of these enhancers in specific cellular or developmental contexts, but that do not drive activity in AML cells. Additional studies will be necessary to understand whether hypermethylation of these enhancer elements is a consequence of decreased occupancy of these modulating factors^47^, or whether it directly prevents proper regulation in ways that contribute to AML development.

## Acknowledgements

This work was supported by the National Cancer Institute (K08CA190815) and The Cancer Research Foundation Young Investigator Award to D.H.S. Support for procurement, annotation, and sequencing of human samples was provided by the Genomics of AML Program Project (P01CA101937, to Dr. Ley) and the Specialized Program of Research Excellence in AML (P50CA171963, to Dr. Link) from the NCI. Sequencing was performed by the McDonnell Genome Institute at Washington University in St. Louis.

