## Supplemental Figures for "Genome-wide analysis of focal DNA hypermethylation in *IDH*-mutant AML samples"

**Figure S1.** *IDH*$^{mut}$ DMRs (AML vs. CD34+ cells) share similar genomic features with other AML subtype DMRs and exhibit low steady state methylation in normal CD34+ cells. A. Percent overlap of AML subtype DMRs with defined genomic annotations. B. Distribution of mean methylation values across *IDH*$^{mut}$ hypermethylated DMRs in *IDH*$^{mut}$ AML samples vs. CD34+ HSPCs. C. Distribution of mean methylation values across *IDH*$^{mut}$ hypermethylated DMRs in *IDH*$^{mut}$ AML samples vs. normal myeloid cells (n=3 promyelocyte samples; n=3 polymorphonuclear leukocyte samples, n=2 monocyte samples).

**Figure S2.** *TET2*$^{mut}$-associated hypermethylation is distinct from canonical CpG-island hypermethylation and is consistent with regions of increased TET2 hydroxymethylation activity in *TET2*$^{wt}$ cells**.** A. Percent overlap of generic AML-associated hypermethylation and *TET2*$^{mut}$-specific DMRs with defined genomic annotations. B. Distribution of CpG density across the set of commonly hypermethylated CpG islands and *TET2*$^{mut}$- specific DMRs. C. CpG conversion rate of 4 paired whole-genome bisulfite and oxidative bisulfite prepared libraries (n=1 for *TET2*$^{wt}$; n=3 for *TET2*$^{mut}$). D. Genome-wide average 5hmc levels across ~10.6 million CpGs with > 10x coverage in each of the paired samples, as calculated by subtracting oxidative bisulfite levels from bisulfite levels. E. Example locus encompassing the *ETV6* gene with diminished 5hmc levels in three *TET2*$^{mut}$ samples (red tracks) compared with a *TET2*$^{wt}$ sample (blue track). F. Distribution of estimated 5hmc levels across 4008 *IDH*$^{mut}$ – specific DMRs, 4586 commonly hypermethylated regions in AML, and 4650 heterochromatic regions in a representative *TET2*$^{mut}$ sample and the *TET2*$^{wt}$ sample.

**Figure S3.** Individual samples with *IDH* mutations alone and in combination with *DNMT3A-R882* exhibit group level methylation trends at *IDH*$^{mut}$-specific and *DNMT3A-R882* DMRs. A. Methylation values across *IDH*$^{mut}$-specific DMRs in a set of 15 *IDH*$^{mut}$ samples (red underline) and 7 *DNMT3A*$^{R882}$/*IDH* doubly mutant samples (blue underline) assayed with WGBS. B.

Methylation values across *IDH*$^{mut}$-specific DMRs in a set of 20 *IDH*$^{mut}$ samples (red underline) and 6 *DNMT3A*$^{R882}$/*IDH* double mutant samples (blue underline) assayed with methylation array. C. Methylation value across *DNMT3A*$^{R882}$ DMRs in a set of 6 *DNMT3A*$^{R882}$ samples (red underline) and 7 *DNMT3A*$^{R882}$/*IDH* doubly mutant samples (blue underline) assayed with WGBS. D. Methylation values across *DNMT3A*$^{R882}$ DMRs in a set of 18 *DNMT3A*$^{R882}$ samples (red underline) and 6 *DNMT3A*$^{R882}$/*IDH* double mutant samples (blue underline) assayed with methylation arrays. E. Hierarchical clustering of CpG methylation values contained within 2183 *IDH*$^{mut}$-specific DMRs in primary AML samples with *IDH1* (n=7), *IDH2* (n=13), *TET2* (n=15), and *DNMT3A*$^{R882}$ (n=6), and co-occurring *DNMT3A*$^{R882}$/*IDH* (n=6), and also *MLL-ELL* (n=11), *CBFB-MYH11* (n=12), and *RUNX1-RUNX1T1* (n=7) fusions. F. Hierarchical clustering of CpG methylation values contained within 3852 *DNMT3A*$^{R882}$ DMRs in primary AML samples with *IDH1* (n=7), *IDH2* (n=13), *TET2* (n=15), *DNMT3A*$^{R882}$ (n=6), and co-occurring *DNMT3A*$^{R882}$/*IDH* mutations (n=7), and also *MLL-ELL* (n=11), *CBFB-MYH11* (n=12), and *RUNX1-RUNX1T1* (n=7) fusions.

**Figure S4.** ChromHMM states are unique in subtype-specific DMRs for AMLs with canonical fusions and *DNMT3*A$^{R882}$ mutations compared to *IDH*$^{mut}$ AML. A. Percent overlap of 1921 RUNX1-RUNX1T1 DMRs with 15 ChromHMM chromatin states. B. Percent overlap of 276 *MLL-ELL* DMRs with 15 ChromHMM chromatin states. C. Percent overlap of 309 *CBFB-MYH11* DMRs with 15 ChromHMM chromatin states.

**Figure S5.** *IDH*$^{mut}$-specific enhancer DMRs are enriched in 'superenhancers' and contact highly expressed genes in AML. A. Locus heatmap of mean subtype methylation across *IDH*$^{mut}$-specific DMRs, including annotated overlaps with gene promoters (green), putative active enhancers (purple), and FitHiC loop anchors (blue). B. Representative rank-ordered analysis of H3K27ac marked enhancers in two *IDH*$^{wt}$ AML samples annotated by enhancer and superenhancer overlap with *IDH*$^{mut}$-specific DMRs. C. Distribution of number of *IDH*$^{mut}$-specific DMRs overlapping computationally defined 'superenhancers' in 3 *IDH*$^{mut}$ AML samples. D. Hierarchical clustering of *IDH*$^{mut}$ -eDMR target gene expression in *IDH1* (n=6), *IDH2* (n=14), and normal CD34+ cord blood cells (N=17, GSE48846). E. Example *IDH*$^{mut}$ -eDMR locus displaying robust interactions with the *DOT1L* promoter. A zoomed in view of the locus demonstrates focal enhancer hypermethylation in *IDH1*$^{mut}$ (purple) and *IDH2*$^{mut}$ (green) samples compared with CD34+ cells (blue). Normalized *DOT1L* expression is shown for 17 CD34+ samples, 6 and 14 *IDH1*$^{mut}$ and *IDH2*$^{mut}$ samples, and 91 *IDH*$^{wt}$ samples. F. Example of *IDH*$^{mut}$ -eDMR locus displaying robust interactions with the *SRSF3* promoter. A zoomed in view of the locus demonstrates focal enhancer hypermethylation in *IDH1*$^{mut}$ (purple) and *IDH2*$^{mut}$ (green) samples compared with CD34+ cells (blue). Normalized *SRSF3* expression is shown for 17 CD34+ samples, 6 and 14 *IDH1*$^{mut}$ and *IDH2*$^{mut}$ samples, and 91 *IDH*$^{wt}$ samples.

**Figure S1.**

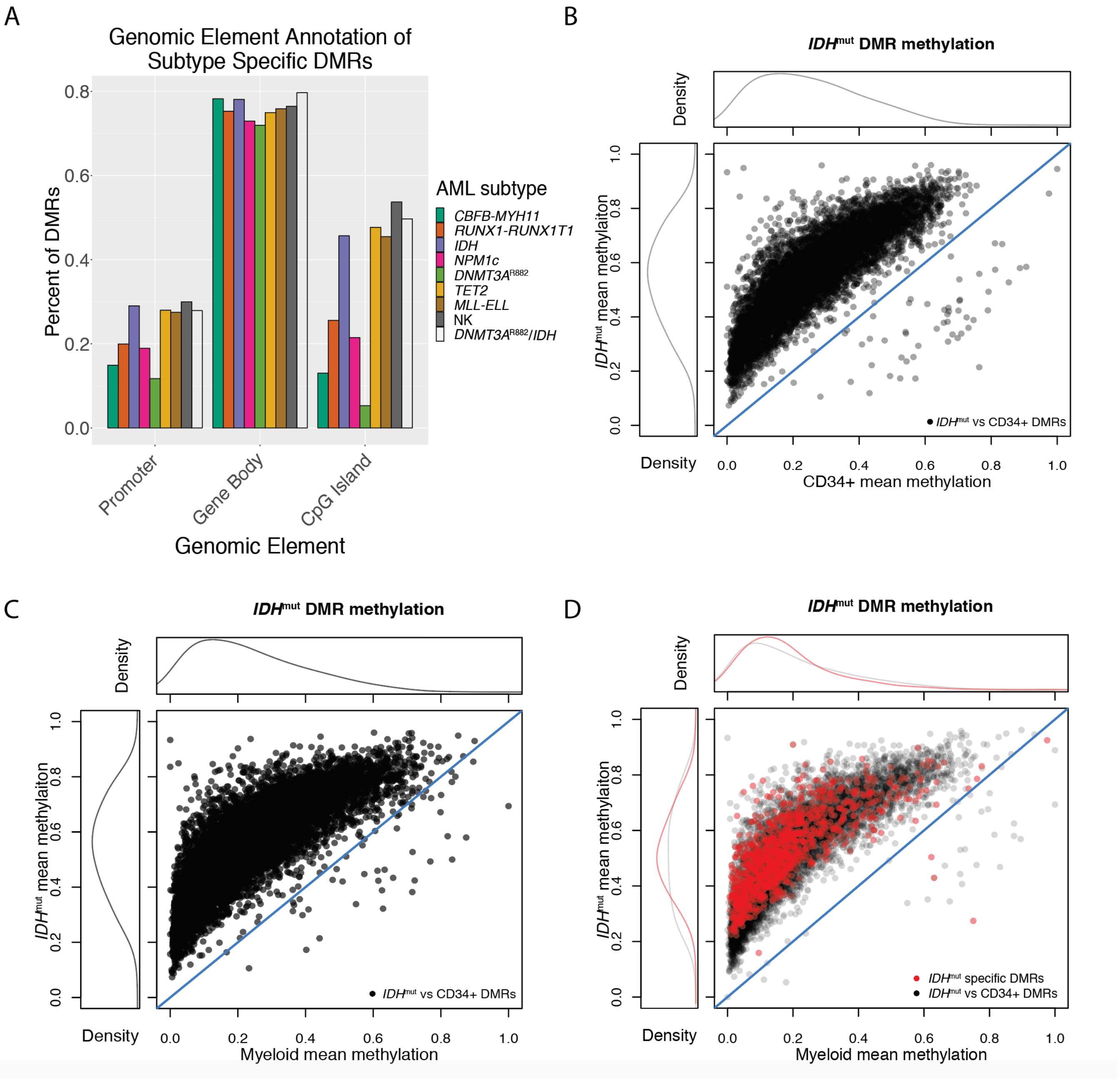
A
Genomic Element Annotation of Subtype Specific DMRs
Percent of DMRs
0.0
0.2
0.4
0.6
0.8
Promoter
Gene Body
CpG Island
Genomic Element
AML subtype
CBFB-MYH11
RUNX1-RUNX1T1
IDH
NPM1c
DNMT3A R882
TET2
MLL-ELL
NK
DNMT3A R882/IDH
B
IDHmut DMR methylation
Density
IDHmut mean methylaiton
IDHmut vs CD34+ DMRs
0.0
0.2
0.4
0.6
0.8
1.0
CD34+ mean methylation
C
IDHmut DMR methylation
Density
IDHmut mean methylaiton
IDHmut vs CD34+ DMRs
Myeloid mean methylation
D
IDHmut DMR methylation
Density
IDHmut mean methylaiton
IDHmut specific DMRs
IDHmut vs CD34+ DMRs
Myeloid mean methylation

**Figure S2.**

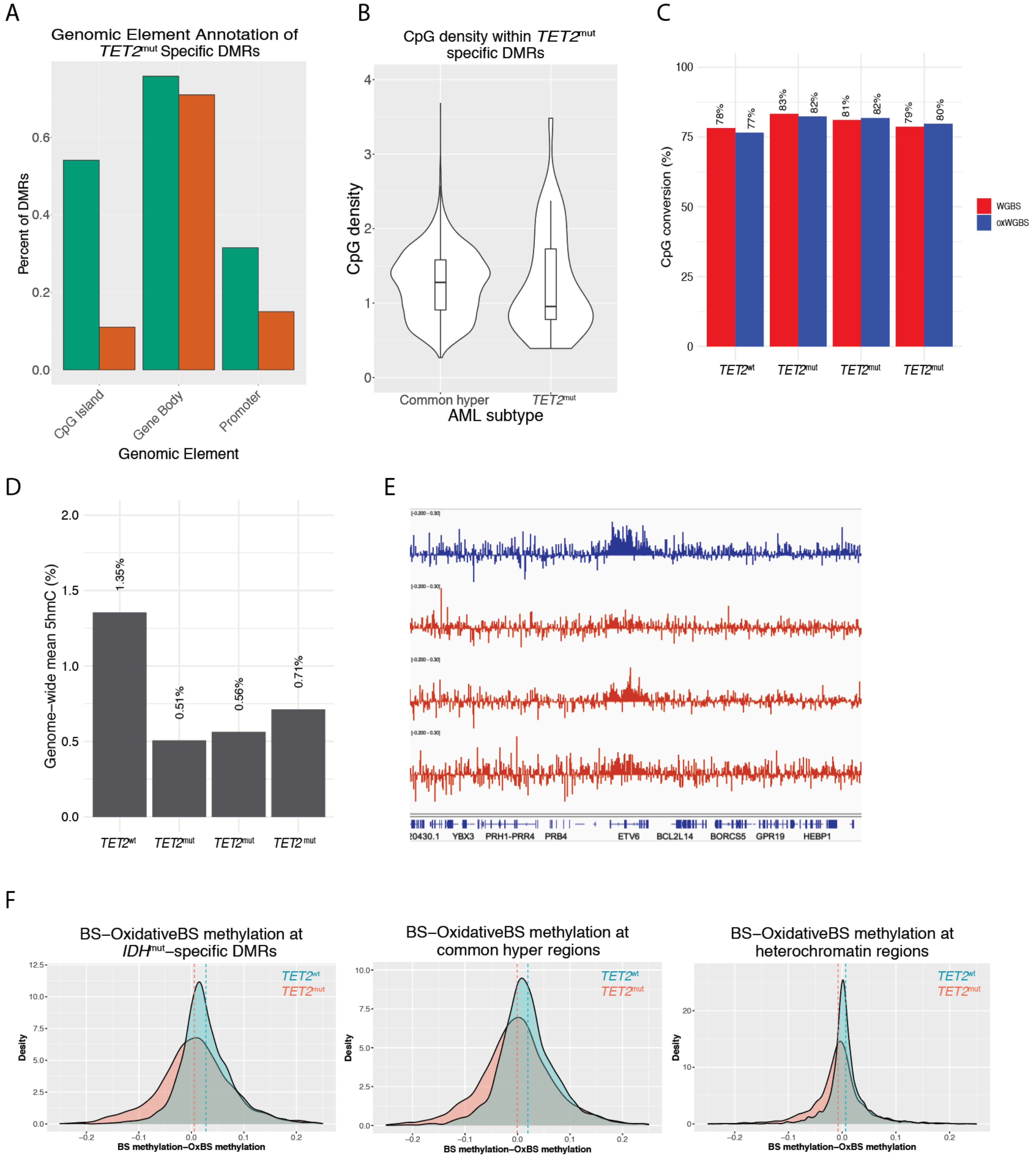
A
Genomic Element Annotation of TET2mut Specific DMRs
Percent of DMRs
0.0
0.2
0.4
0.6
CpG Island
Gene Body
Promoter
Genomic Element
B
CpG density within TET2mut specific DMRs
CpG density
0
1
2
3
4
Common hyper
TET2mut
AML subtype
C
CpG conversion (%)
0
25
50
75
100
78%
77%
83%
82%
81%
82%
79%
80%
TET2wt
TET2mut
TET2mut
TET2mut
WGBS
oxWGBS
D
Genome-wide mean 5hmC (%)
0.0
0.5
1.0
1.5
2.0
1.35%
0.51%
0.56%
0.71%
TET2wt
TET2mut
TET2mut
TET2mut
E
YBX3
PRH1-PRR4
PRB4
ETV6
BCL2L14
BORCS5
GPR19
HEBP1
F
BS−OxidativeBS methylation at IDHmut−specific DMRs
BS−OxidativeBS methylation at common hyper regions
BS−OxidativeBS methylation at heterochromatin regions
TET2wt
TET2mut
Desity
BS methylation−OxBS methylation

### Figure S3.

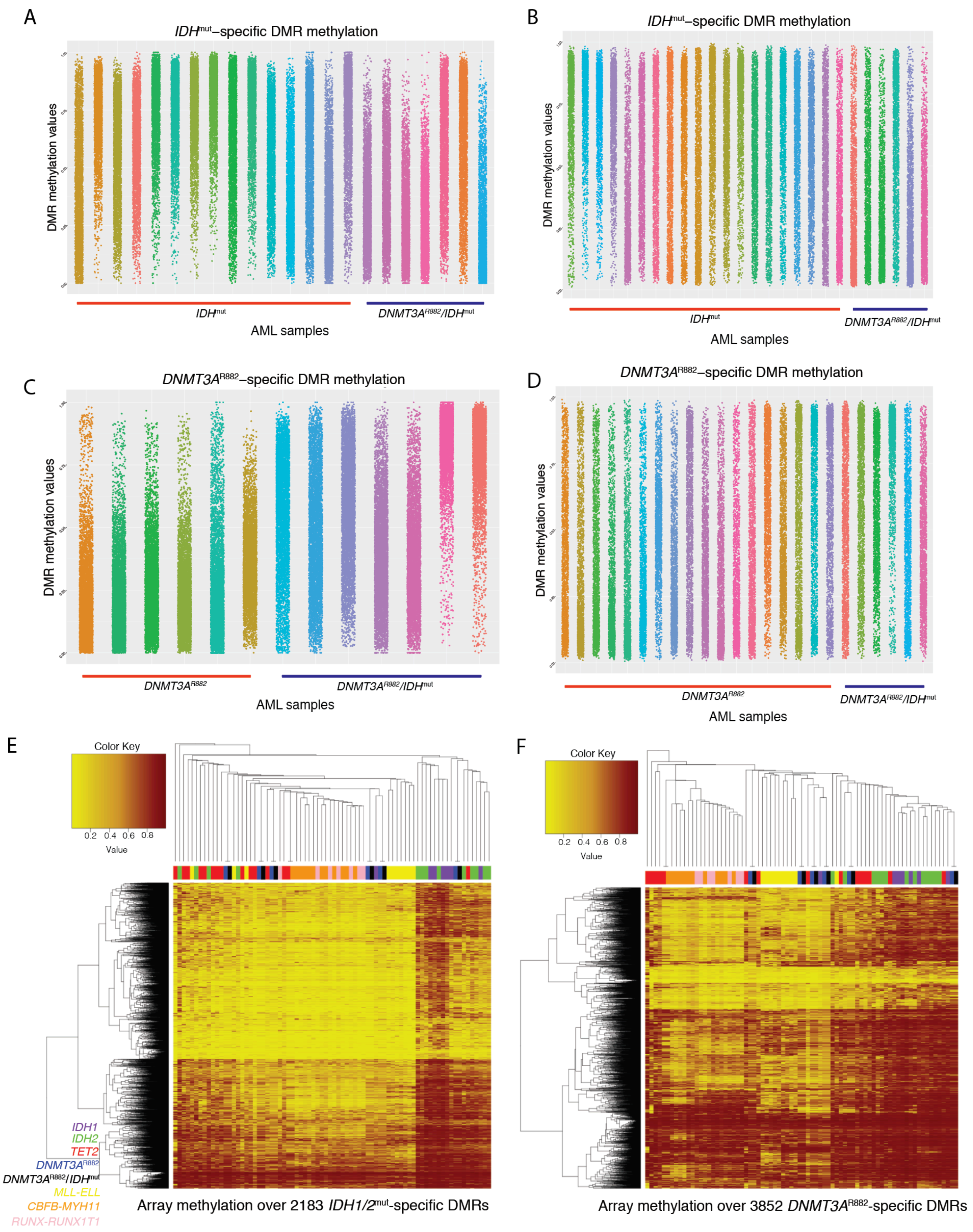

A
IDHmut–specific DMR methylation
DMR methylation values
IDHmut
DNMT3AR882/IDHmut
AML samples
B
IDHmut–specific DMR methylation
DMR methylation values
IDHmut
DNMT3AR882/IDHmut
AML samples
C
DNMT3AR882–specific DMR methylation
DMR methylation values
DNMT3AR882
DNMT3AR882/IDHmut
AML samples
D
DNMT3AR882–specific DMR methylation
DMR methylation values
DNMT3AR882
DNMT3AR882/IDHmut
AML samples
E
Color Key
0.2 0.4 0.6 0.8
Value
IDH1
IDH2
TET2
DNMT3AR882
DNMT3AR882/IDHmut
MLL-ELL
CBFB-MYH11
RUNX-RUNX1T1
Array methylation over 2183 IDH1/2mut-specific DMRs
F
Color Key
0.2 0.4 0.6 0.8
Value
Array methylation over 3852 DNMT3AR882-specific DMRs

**Figure S4.**

A

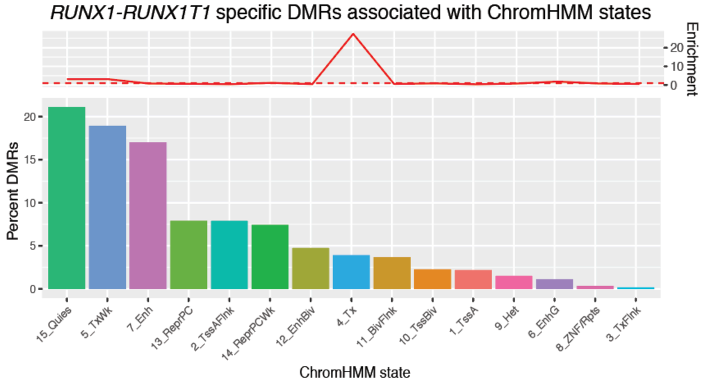


B

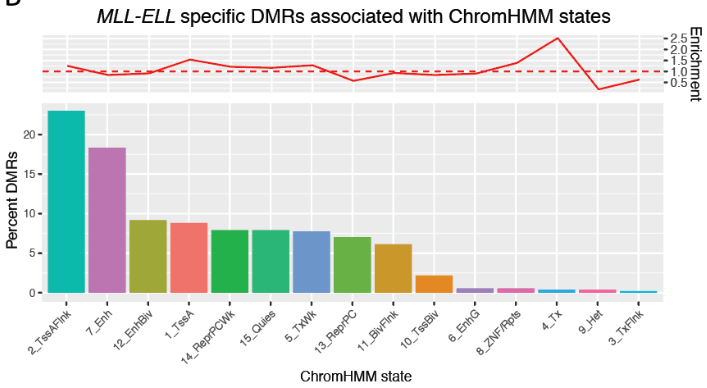


C

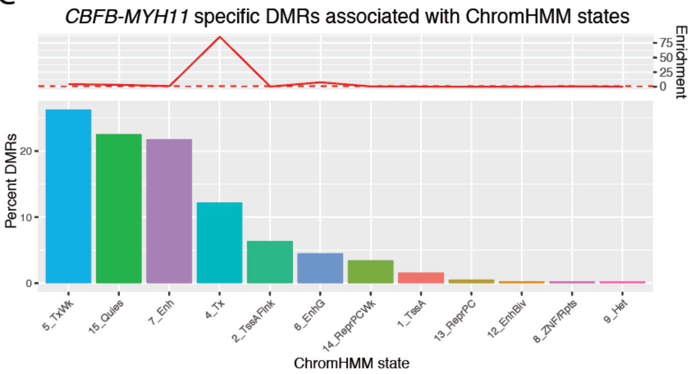

**Figure S5.**

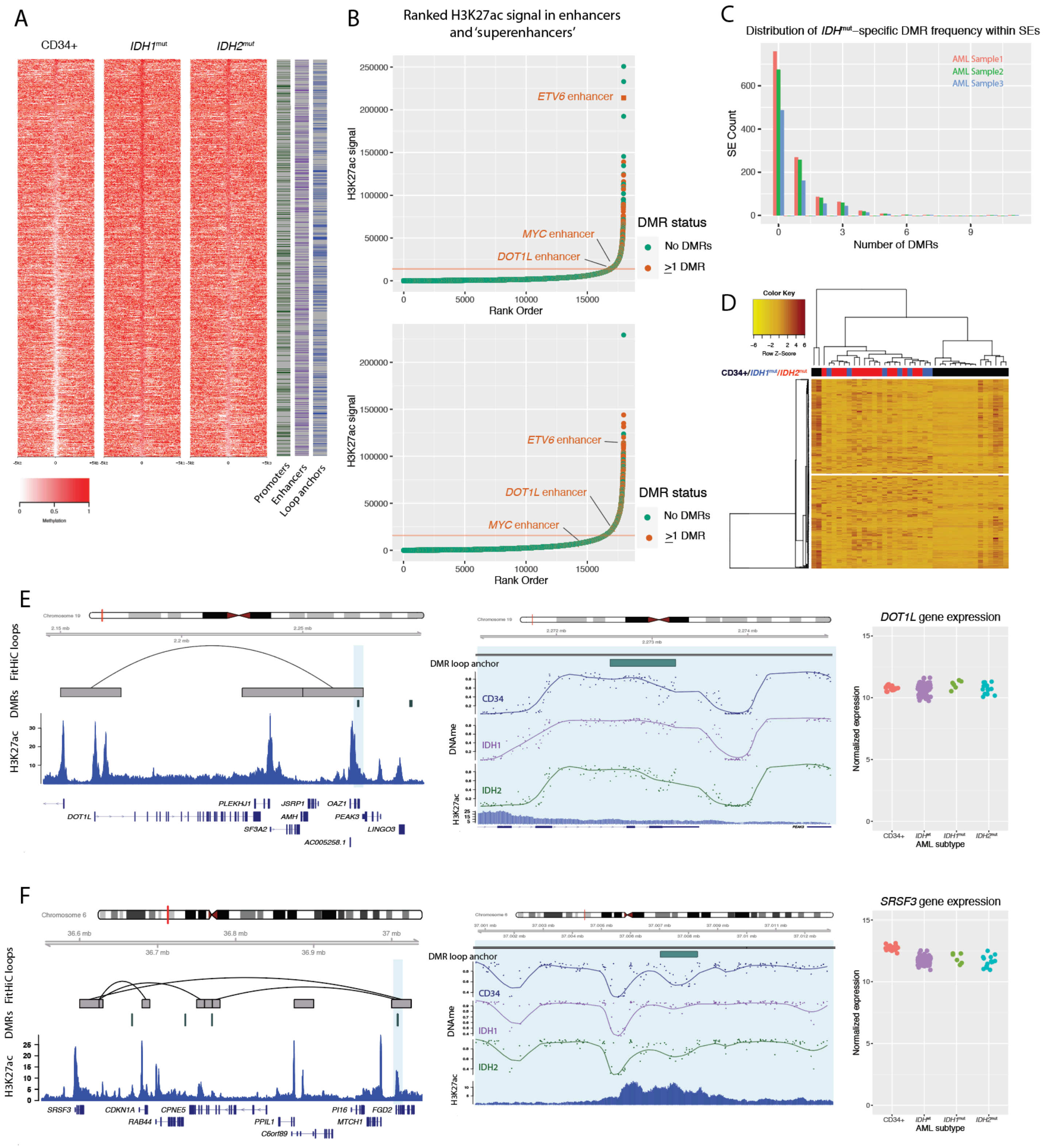

A
CD34+
IDH1mut
IDH2mut
Promoters
Enhancers
Loop anchors
Methylation
B
Ranked H3K27ac signal in enhancers and 'superenhancers'
H3K27ac signal
Rank Order
ETV6 enhancer
MYC enhancer
DOT1L enhancer
DMR status
No DMRs
>1 DMR
C
Distribution of IDHmut-specific DMR frequency within SEs
AML Sample1
AML Sample2
AML Sample3
SE Count
Number of DMRs
D
Color Key
Row Z-Score
CD34+/IDH1mut/IDH2mut
E
FitHiC loops
DMRs
H3K27ac
DMR loop anchor
CD34
IDH1
IDH2
DNAme
DOT1L gene expression
Normalized expression
AML subtype
F
SRSF3 gene expression
